# Methane efflux from desert soils

**DOI:** 10.1101/2025.09.19.677267

**Authors:** Ilya Gelfand, Vasily I. Grabovsky

## Abstract

Deserts cover ~50% of Earth’s surface but are poorly studied regarding their methane (CH_4_) fluxes. Current theory suggests that dry soils act as CH_4_ sinks, with higher soil carbon (C) content enhancing oxidation. To test this, we measured CH_4_ fluxes in 12 field plots with increasing C content in Israel’s Negev Desert over three summer and three winter months using automated chambers. We hypothesized that well-aerated soils would oxidize CH_4_, and the oxidation rate will increase with more organic matter.

Contrary to our hypothesis, CH_4_ was emitted ~75% of the time, with average effluxes of 0.222 ± 0.007 nmol m^−2^ s^−1^ in summer and 0.081 ± 0.004 in winter. We propose that CH_4_ emissions were driven by abiotic processes related to solar radiation, not microbial activity. A shading experiment reduced CH_4_ efflux by 51%, supporting this idea.

The newly identified CH_4_ source of at least 89.8 ± 2.2 mg CH_4_ m^−2^ yr^−1^ equals ~30% of the modelled CH_4_ sink (~300 mg CH_4_ m^−2^ yr^−1^) for the same region.

**Synopsis:** Our field measurements in Israel’s Negev Desert show that dry desert soils can emit methane, challenging current models that assume these soils consistently act as atmospheric methane sinks.

## Introduction

Methane (CH_4_) is the second most important greenhouse gas (GHG) with a global warming potential (GWP) ~28 times that of carbon dioxide (CO_2_). CH_4_ contributes 15%– 35% to the additional radiative forcing from anthropogenic emissions of GHGs and their precursors since the industrial revolution ^1^. Current global CH_4_ emissions are estimated at 540–865 Tg CH_4_ yr^−1^, while total sinks are estimated at 507–796 Tg CH_4_ yr^−1^. The present atmospheric CH_4_ concentration is ~2 ppmv (µL L^−1^), and in 2022 the rate of increase was ~14 ppbv yr^−1^ (nL L^−1^ yr^−1^), or ~38 Tg CH_4_ ^2,3^.

Natural sources, primarily wetlands and other anaerobic environments, account for ~45% of global CH_4_ emissions, with the remainder from anthropogenic sources such as agriculture and fossil fuel use ^3,4^. In soils, CH_4_ is produced by specialized microbes in anaerobic environments, while CH_4_ production in dry soils is considered negligible ^5,6^ because no microorganisms are known to produce CH_4_ under aerobic conditions ^7^. However, over the past ~20 years, multiple studies have reported potential abiotic CH4 sources from soils under controlled conditions ^8–11^, including photochemical cleavage of ketones ^12^, methyl radicals, and hydroxyl radical reactions ^13^.

The primary global sinks of CH_4_ are soil oxidation (~6%) and photochemical reactions in the troposphere (~90%). While the soil sink is smaller, it remains important because it is comparable to the annual increase in atmospheric CH_4_ due to anthropogenic emissions (11– 49 vs. 20 – 43 Tg CH_4_ yr^−1^) ^2,3,14^. Soil CH_4_ oxidation is mediated by methanotrophic microbes and influenced by factors such as soil oxygenation (i.e., water content and gas diffusivity), CH_4_ concentration within the soil profile, and methanotroph diversity ^7,15,16^. In contrast to ammonium (NH_4_^+^) and nutrient availability ^16^, tillage (i.e., soil disturbance) does not affect soil CH_4_ oxidation rates ^17,18^. Increasing soil organic carbon (SOC) content, enhances CH_4_ oxidation, particularly in upland forest soils, likely due to improved soil structure (e.g., aggregation), oxygenation, and microbial diversity ^19^.

Understanding and quantifying the global CH_4_ budget is crucial for developing realistic climate change mitigation strategies ^4,20^. The current imbalance of 15–38 Tg CH_4_ yr^−1^ is largely due to poorly constrained soil sources and sinks ^3^ driven by limited spatial measurement coverage in terrestrial ecosystems ^21^ and a scarcity of data from arid environments ^22,23^. Deserts cover ~50% of Earth’s land area ^24^ and are thought to be a significant CH4 sink, currently estimated at 4.9 Tg CH_4_ yr^−1 25^. However, due to limited field measurements, this desert sink remains poorly constrained in global CH_4_ budgets and models ^26–29^.

Here, we present the first comprehensive in situ assessment of CH_4_ fluxes from desert soils in the Middle East. We used automated chambers in the Negev Desert, with experimental manipulation of SOC levels, to estimate the potential contribution of arid soils to the CH4 sink. Six replicated chambers were deployed in an open field for three summer and three winter months, measuring 10,410 individual CH_4_ fluxes from soils with three SOC levels, each replicated four times. We hypothesized that dry desert soils would oxidize CH_4_, with oxidation rates increasing alongside SOC content.

## Materials and Methods

### Site description

The study was conducted at the Ben Gurion University of the Negev, Sde Boker campus (30°51′08.27″N, 34°47′00.24″E; 480 m above sea level), for a ~3 months in the summer (May – July) and ~3 months during the winter (November – March). The Negev Desert has an arid Mediterranean climate, with hot rainless summers and cool wet winters. Precipitation mainly occurs during October–April with the rest of the year is dry. The air temperature during the study period ranged between 2.8 and 40.5 °C with an average of 25.6 °C during the summer and 13.5 °C during the winter months. The summer was dryer than the winter with average daily RH 50.1% during the summer and 66.6% during the winter. The average solar radiation load during the summer months was 25.9 ± 3.2 MJ m^−2^ day^−1^ and during the winter months 12.4 ± 2.9 MJ m^−2^ day^−1^, while the soil temperature (0–10 cm depth) was between 21.6 and 41.5 °C and precipitation was 21.4 mm during the winter months (Figure S1).

### Study area

The area selected for the field experiment was bare. Four years prior to the start of measurements, we set three experimental treatments in four replicate areas that were randomly distributed across the field area: tilled soil (rototilled to a depth of 20 cm); tilled soil with addition of 2.6 kg m^−2^ of dried and ground maize stalks (C/N ratio of 57); and tilled soil with addition of 5.1 kg m^−2^ of dried and ground maize stalks. Each replicate area was ~1.8 m wide and ~15 m long, with 0.5 to 1 m-wide passages between the replicate areas. In addition to these passages, the replicated areas were demarcated along their long sides by 2 mm-thick PVC sheets buried to a depth of ~one m to prevent lateral water flow and material transfer. Three months before the start of the measurements all replicates were tilled to a depth of ~20 cm and raked to homogenize the soil.

The soil in the study area is a deep sandy loam loess, with 59% sand, 30% silt, and 11% clay and without any apparent horizons or aquifer, at least in its upper 60 m ^30^. The bulk density of the upper 20 cm of soil is 1.24 ± 0.04 g cm^−3^, and the water-holding capacity (WHC; sieved soil) is 44.5% ± 0.9%. The soil at the field site (0 – 20 cm depth) before the addition of the ground maize contained 0.22 ± 0.12 wt.% C and 0.05±0.03 wt.% N (mean and standard error; *n* = 4 composite samples), and the pH was 8.24±0.03 (1:5 w:v; soil:water).

### Automated chamber and auxiliary measurements

Surface fluxes of CH_4_ were measured continuously between 2 May 2023 and 27 July 2023 and between 29 November 2023 and 3 March 2024, using an automated chamber system installed in the field ~4 weeks before the start of the measurements. The system consisted of seven opaque, automated gas flux chambers connected to a multiplexer (eosAC and eosMX; Eosense Inc. Dartmouth, Nova Scotia, Canada). The multiplexer allows for dynamically signaled chamber deployment and routed the gases to an off-axis integrated cavity output spectroscope (OA-ICOS) capable of measuring CH_4_ concentrations in the field with a precision of 0.001 ppmv (µL L^−1^) (GLA331N2OM1; ABB, Quebec City, Canada). The chambers were measured sequentially over a 5-min sampling period with a 5-min flushing period before and after each measurement. Six chambers were deployed linearly along replicate areas with ~2 m between the two closest chambers. The chambers were moved between the replicates and different treatments every 2–5 days to cover all areas on a ~3×2 m grid.

A dummy seventh chamber was deployed for measurement of erroneous soil fluxes in order to estimate minimum detectable flux (MDF) based on the precision of our instrumentation and overall measuring system uncertainty (see SI for details). The dummy chamber was deployed on a metal sheet, had no direct contact with the soil, and stayed in place for the duration of the experiment. Chamber collars were installed in all the replicates during the measurements, and their height was measured each time the chambers were moved to ensure that the chamber volumes did not change over time. During the 21 days of June chambers were removed from the field and set on the lifted steel mesh platform that allowed free air movement in and out of the chambers during closure times. This was done to assess overall system performance and rule out erroneous changes in chamber headspaces gas concentrations that may affect soil gas flux measurements.

Soil organic C contents were measured in 10 soil samples from the 0–20 cm depth that were collected with a hand auger (2.5 cm internal diameter) from each replicate area. Soil C content was measured in soil that passed through a 500 µm sieve by combustion in a CHNS/O analyzer (FlashSmart™ Elemental Analyzer; Thermo Fisher Scientific) after carbonate removal by oxidation with HCl ^31^. The air temperature and pressure inside the chambers were measured with sensors installed in the chambers by the manufacturer.

### Solar irradiation exclusion experiment

To assess the effect of solar radiation on soil CH_4_ flux, we conducted an exclusion experiment over three weeks in June 2025. A one m^2^ frame of ~ 1.5 m height was installed above each of the three automated chambers and covered with reflective film to block direct sunlight. An additional three automated chambers, located approximately two meters away, remained uncovered and served as control.

### Limit of detection estimation

We calculated a minimum detectable slope (MDS) of ±0.0045 and ±0.0055 ppbv CH_4_ s^−1^ according to the statistical approach of ^32^ using the CH_4_ concentration measurements from all incubations of the chamber deployed on a metal sheet without soil contact (*n* = 664 chamber incubations and 17928 individual concentration values for three summer months and *n* = 1120 chamber incubations and 30240 individual concentration values for three winter months; see the SI text for details). For the estimation of soil CH_4_ emissions we excluded the data with slopes less than the MDS (i.e., slopes of <0.0045 and 0.0055 ppbv CH_4_ s^−1^; Table S2–S3). After removal of the MDS, we estimated minimum detectable fluxes (MDF) of 0.1188±0.005 and 0.0584 ± 0.0002 nmol CH_4_ m^−2^ s^−1^ for summer and winter months, respectively. The MDF was estimated as the average of erroneous soil CH_4_ “emissions” measured by the dummy chamber deployed on the metal sheet without soil contact (see the SI text for details). We then subtracted the MDF from the reported fluxes.

### Data analysis

Flux calculations and analyses were performed in R ^33^, as were all statistical analyses used for the flux calculations. Specifically, we used R packages (dgof, fitdistrplus ggplot2, readxl, writexl, dygraphs, reshape, dplyr, tidyverse, anytime, sjmisc, lubridate, stats, tsibble, imputeTS). The normality of the residuals was assessed by examining histograms and normal probability plots. The homogeneity of the residual variances was examined with the Shapiro–Wilk and Levene’s tests. When the normal probability plots suggested the residual distribution deviated from normality, the Wilcoxon nonparametric test was applied to the data analysis. Multiple comparisons amongst the treatments were analyzed using a mixed model approach ^34^. The statistical models for the studied data consisted of the fixed effect of the soil C content. The models also included the random effects of the replicate (n = 4 per C level) and gas measurement chamber (n = 6) interactions. The chambers were nested within replicates of different C levels. Normality of the residuals was assessed by examining histograms and normal probability plots. The homogeneity of residual variances was assessed by Levene’s test. When normal probability plots suggested that the distributions of the residuals deviated from normality, the response variables were transformed as needed to conform with normal distribution. Whenever the violations of the variance homogeneity assumption were not fully addressed by the transformations, unequal variance analyses were performed following the approach of ^34^. Results of statistical analyses are presented in the supplemental materials. In order to make the data readily comparable with other published sources, we only reported the observed data, their mean values, and standard errors in the original scale, even when the statistical analyses were conducted using transformed data as described above. We calculated the diurnal soil CH_4_ emissions by averaging all measured soil CH_4_ fluxes hourly during rainless and hot summer and wet and cool winter months. The mean fluxes for summer and winter were calculated by linear interpolation between individually measured during the experiment fluxes by each chamber ^35,36^. Total dataset included 3904 and 6444 individual flux measurements for summer and winter months, respectively.

### Time series analysis

To extract diurnal changes in measured fluxes we used (decompose) R function, a technique used to break down a time series into its constituent components: trend, seasonality, and residuals (noise). We converted the raw data with function ts() using 24-hour frequency and filled gaps in the raw data using has_gaps(), scan_gaps() И count_gaps() from the tsibble library. The gaps were filled using na_locf() function from imputeTS library.

### Assessing global implications

To assess the potential importance of measured CH_4_ fluxes from the desert soils, we used average fluxes for summer and winter months, respectively. Further we identified global distribution of soils and climatic conditions similar to climate and soils at our site using Google Earth Engine (GEE). Parameters included in the GEE are provided in Table S4. Currently, the global CH_4_ observations dataset does not include measurements in sites similar to the Negev Desert and/or Middle East climate and soil type ^25^.

## Results and Discussion

### CH_4_ fluxes from soil

Soil CH_4_ fluxes during the summer months were mostly positive and increased linearly with topsoil organic matter content, ranging from 0.179 ± 0.008 to 0.293 ± 0.014 nmol CH_4_ m^−2^ s^−1^ (mean ± standard error after removal of the minimum detectable flux; n = 1075–1547 [individual measured fluxes per SOC content]; Fig. 1a). During winter months, CH4 fluxes were also mostly positive and related to SOC levels, though lower— between 0.040 ± 0.006 and 0.145 ± 0.007 nmol CH_4_ m^−2^ s^−1^ (mean ± standard error after removal of the minimum detectable flux; n = 2026–2342 [individual measured fluxes per SOC content]; Fig. 1b). Increased SOC content had little effect on diurnal CH_4_ flux patterns. CH4 fluxes showed pronounced diurnal variability: emissions were higher during the day and closer to zero at night (Fig. 2). In both summer and winter, ~12% of CH_4_ fluxes were negative, indicating oxidation, though net efflux dominated. Beyond diurnal and seasonal patterns, fluxes were highly skewed and variable (Fig. S2; Table S1), indicating strong spatial heterogeneity—common in trace gas fluxes measured by static chambers ^37^.

**Figure 1.**
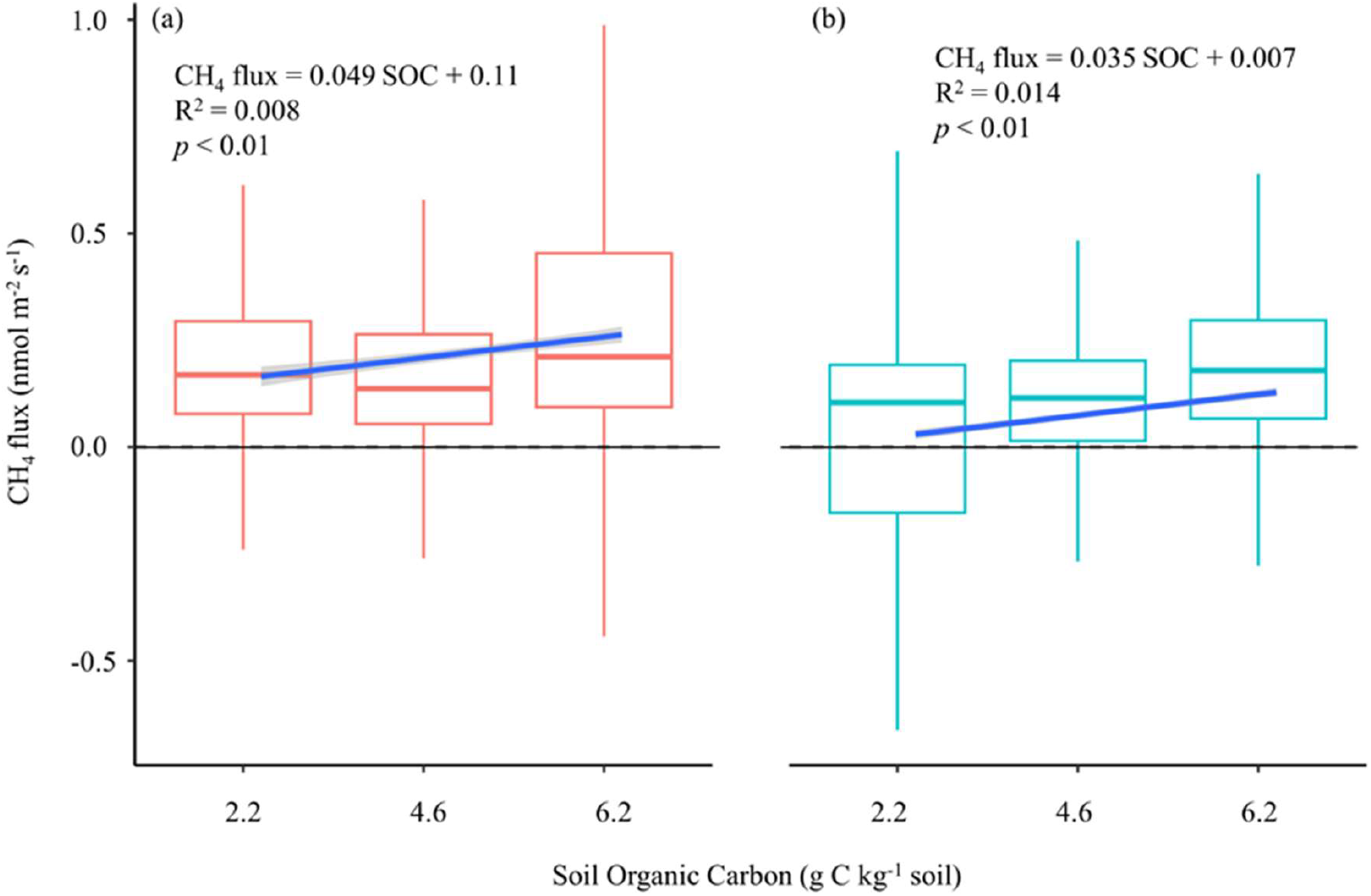
Soil CH_4_ flux (nmol m^−2^ s^−1^) measured during summer (a) and winter (b) months from soils with increasing organic carbon content (g C kg^−1^ soil). Boxplots represent median, 25^th^ and 75^th^ percentiles and minimum and maximum values. See Table S5 for statistical analysis of soil carbon content effect on CH_4_ fluxes during summer and winter months.

**Figure 2.**
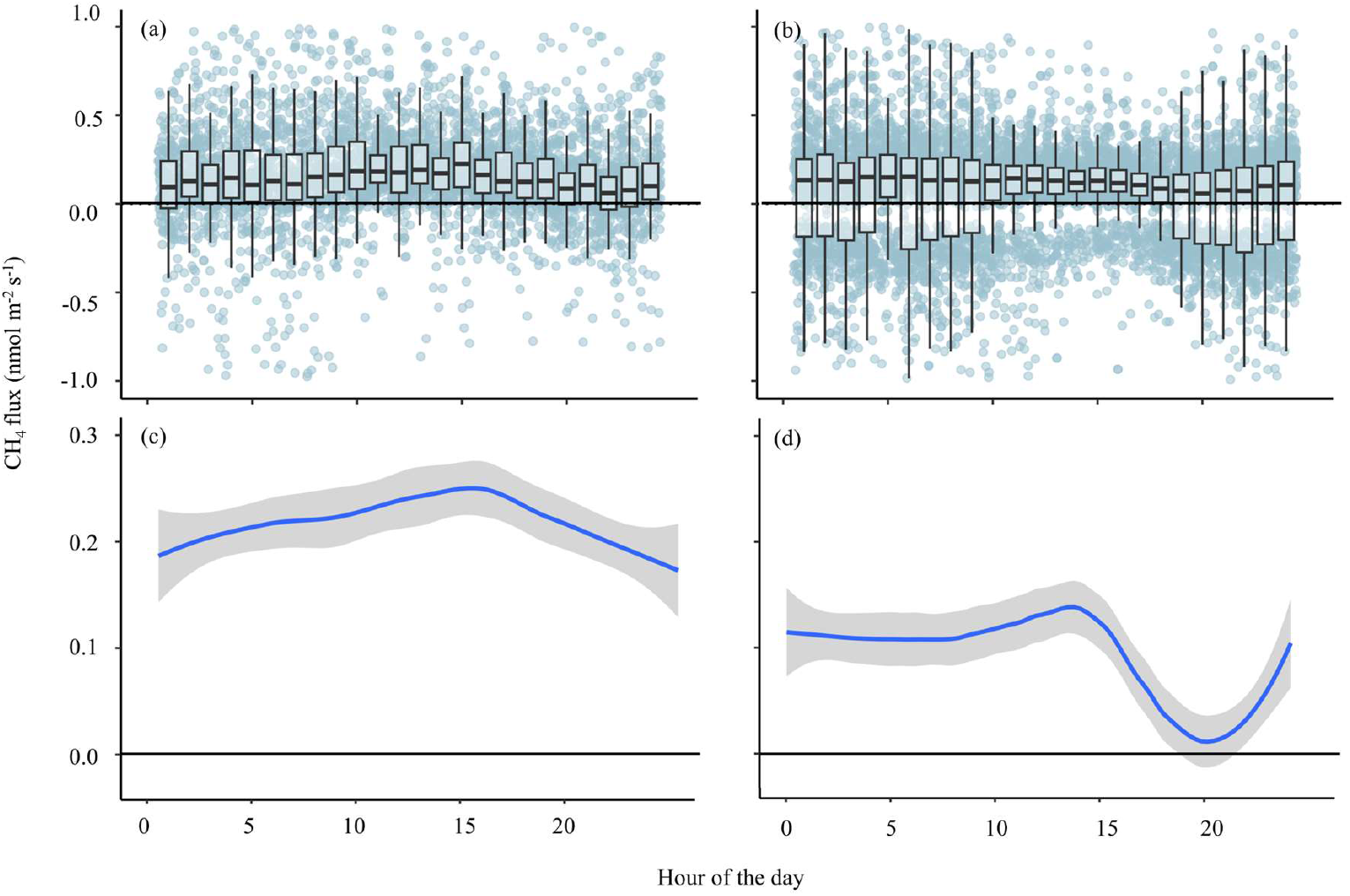
Diurnal dynamics of soil CH_4_ flux (nmol m^−2^ s^−1^) calculated by averaging all fluxes measured every 15 min within a 1 h period in summer (a) and winter (b) months; (d) and (c) present diurnal flux dynamics after removal of random component by time-series analysis. Boxplots represent median, 25^th^ and 75^th^ percentiles and minimum and maximum values of all fluxes measured within a given hour (130–193 individual fluxes h^−1^). Background dots on (a) and (b) panels show all individual fluxes measured during the field experiment. Attention to y-axis changes across panels.

Our findings contradict the conventional understanding of methanogenesis in drylands ^16,38,39^. Previous studies explored CH_4_ emissions following soil rewetting and hypothesized that anaerobic microsites could form, enabling methanogenesis ^40,41^. Dry soil emissions measured in southeastern Australia ^42^ were attributed to methanogenesis by termites, which are absent in our field. The only CH_4_ data from Negev Desert soils (under biocrusts) did not reveal consistent vertical trends in CH_4_ concentrations ^43^. Aside from that, we found no CH_4_ flux data from the Negev Desert, nor from broader Middle East, Central Asia, or North Africa regions ^17,44^. Overall, desert CH_4_ flux data are scarce; a recent analysis identified only ~30 studies from deserts worldwide ^25^. Most were conducted in cold deserts (annual temperature <10 °C) or regions with summer precipitation.

### Solar irradiation exclusion experiment

Average CH_4_ efflux from sun-exposed chambers (~0.291 ± 0.006 nmol CH_4_ m^−2^ s^−1^; Table S1) was consistent with summer averages. Under sunlight-blocking covers, however, efflux decreased to 0.149 ± 0.005 nmol CH_4_ m^−2^ s^−1^—approximately 50% lower (Fig. 3)—highlighting solar radiation as a driver of CH_4_ emissions.

**Figure 3.**
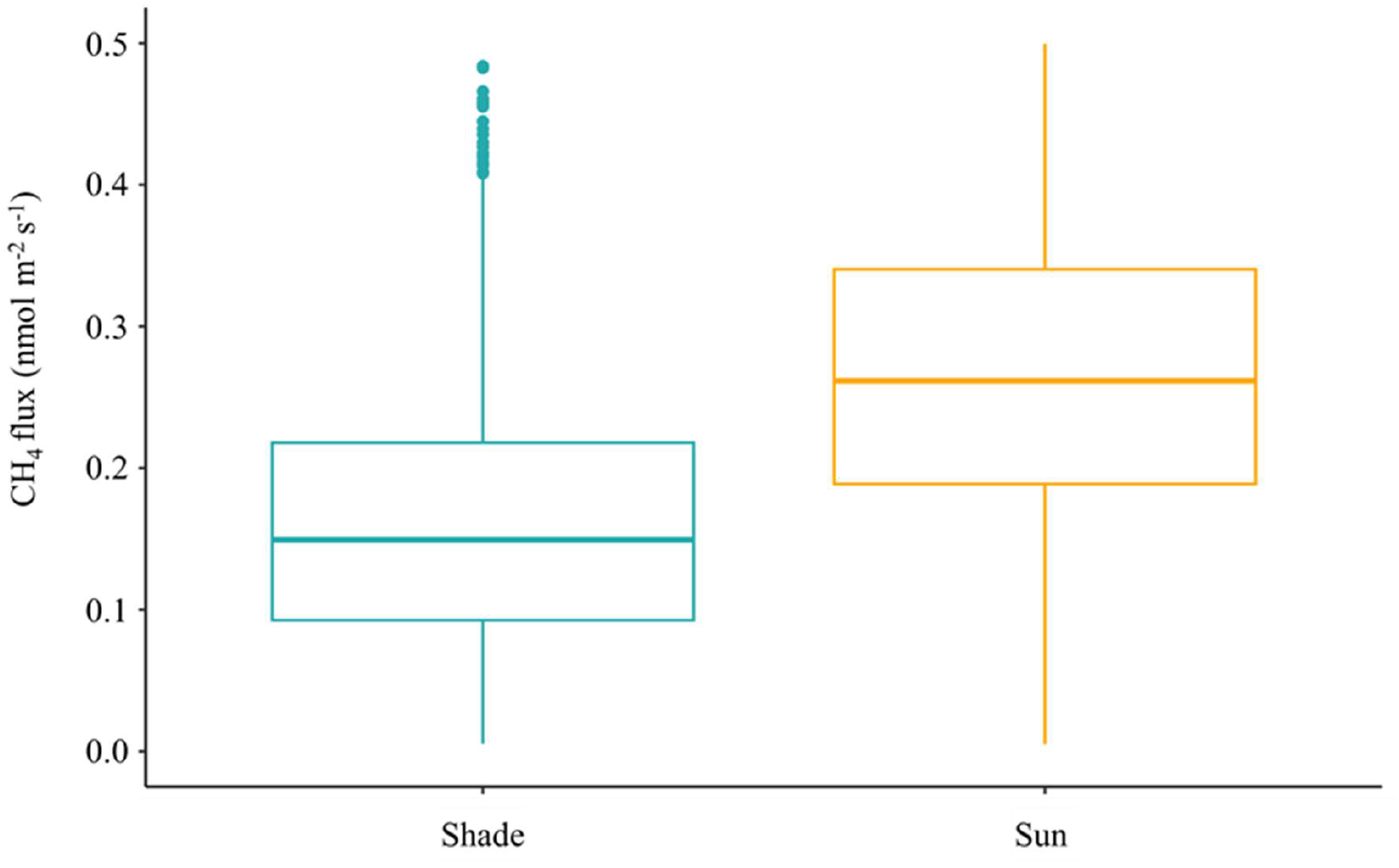
Average soil CH_4_ flux (nmol CH_4_ m^−2^ s^−1^) measured in *Shade*, beneath shelters blocking total direct radiation and in *Sun*, an open area (n = 999 and 994 individual flux measurements, respectively; *p* = 0.0495). Boxplots represent median, 25^th^ and 75^th^ percentiles and minimum and maximum values.

Though data from hot and Mediterranean deserts are limited, CH_4_ emission rates <3 nmol CH4 m^−2^ s^−1^ have been reported from abiotic production mechanisms—namely, photochemical and chemical decomposition of organic matter in aerobic soils ^8–11^. Such abiotic decomposition likely explains both the CH_4_ emissions and the observed correlation with SOC content (Fig. 1). Prior studies also demonstrated abiotic litter breakdown via photodegradation in drylands ^45^ supporting this mechanism. These abiotic reactions merit further experimental validation and field investigation.

Our soils experienced high solar energy loads (>25 MJ m^−2^ day^−1^; Fig. S1) during the dry summer months. In winter, irradiation was nearly halved, and average CH_4_ efflux likewise declined twofold (Table S1). Artificial shading reduced CH_4_ emissions by 51% (Fig. 3), reinforcing the hypothesis that emissions originate from abiotic SOC decomposition.

The presence of negative fluxes suggests multiple interacting processes: photochemical/chemical SOC degradation, methanotrophy, and possible CH_4_ diffusion from deeper soil layers—potentially driven by abiotic gas–water–rock reactions ^46^ from ancient aquifers (>500 m depth) beneath the Negev ^30,47^. CH_4_ concentration in gas extracted from groundwater systems in the Negev (Kurnub and Judea formations) can range from 50 to 5,000 ppmv (Itay Reznik, Geological Survey of Israel, personal communication).

### Global implications

To assess the broader significance of these CH_4_ emissions, we extrapolated average daily fluxes for summer and winter using soil and climate data via Google Earth Engine (GEE). We identified 1,365,000 km^2^ of land globally with conditions similar to our study site (Fig. 4). Using average fluxes of 0.222 ± 0.007 nmol CH_4_ m^−2^ s^−1^ (May–October) and 0.081 ± 0.004 (November–April), we estimate annual emissions of 122.5 ± 3.1 kilotons CH_4_. This represents ~2% of the global soil CH_4_ sink currently attributed to deserts ^25^. Though small, this efflux reverses the assumed role of desert soils from sinks to net sources.

**Figure 4.**
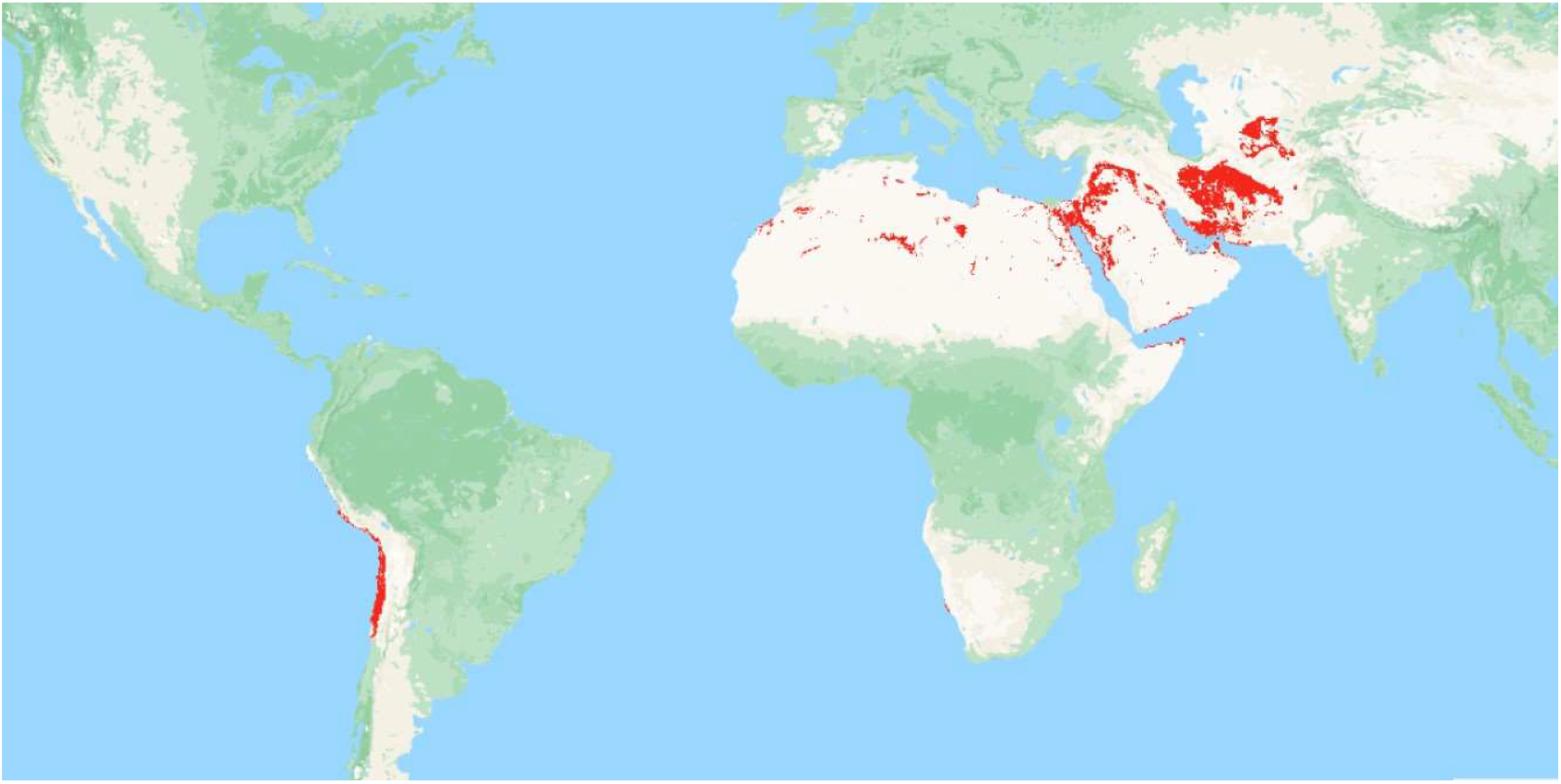
Global distribution of soils (in red) similar to Negev desert with the potential to emit CH_4_. A total area of 1.37 × 10^6^ km^2^ of such area was identified globally.

The newly identified CH_4_ source of at least 89.8 ± 2.2 mg CH_4_ m^−2^ yr^−1^ equals ~30% of the modelled CH_4_ sink (~300 mg CH_4_ m^−2^ yr^−1^) for the same region ^48^. The key discrepancy lies in model assumptions: current CH_4_ models exclude dry soil emissions ^49^. Instead, they assume deserts are neutral or act solely as CH_4_ sinks via oxidation. Our data show that dry desert soils can emit CH_4_—an overlooked process in global CH_4_ budgeting ^27^.

## Supporting information

Detailed methods STables and SFigures

## Acknowledgments

We thank Y. Kuzyakov, S. H. Hamilton, G. P. Robertson, and K.V. Severinov for many helpful comments on an earlier version of this work. Financial support for this work was provided by the Israel Science Foundation (Grant no. 305/20).

## Author Contributions

IG: Conceptualization, Funding acquisition, Project administration, Supervision, Investigation, Methodology, Writing – original draft, writing – review and editing. VIG: Methodology, Investigation, Visualization, Writing – review and editing.

