## Supplementary material for "Methane efflux from desert soils": Detailed methods STables and SFigures

**This file includes:**
SI text
SI Figures S1 and S2
SI Tables S1-S5

**SI text**

**Measurements of soil gaseous flux**

To produce reliable gaseous flux estimates, it is necessary to determine the minimum detectable flux (MDF). An analysis of the MDF of accumulation chamber–GC measurements was presented by (Parkin et al., 2012). With the introduction of QCL-based techniques, the MDF needs to be redefined. The QCL sensitivity and continuous nature of the measurements may allow the measurement of very small soil gaseous fluxes, which may help to constrain the atmospheric budgets of trace gases that are currently not well constrained. There are few estimates of the MDF using QCL-based measurements of soil $CH_4$ flux, (Christiansen et al., 2015) reported uncertainty of 0.019 nmol $m^{-2}$ $s^{-1}$, while Nickerson (2016) estimated MDF of 0.002 nmol $m^{-2}$ $s^{-1}$ (Nickerson, 2016).

We used an array of seven automated soil chambers coupled to an OA-ICOS IRGA with QCL technology for the $CH_4$ concentration measurement to calculate the MDF for soil $CH_4$ fluxes in the soils of the Negev Desert, Israel, during the rainless season between May and July and wet winter months, November – March.

**Estimation of the minimum detectable flux for measurement of soil exchange of $CH_4$ with automatic chambers coupled to infrared gas analyzers**

Two main factors affect the measurement uncertainties of soil–atmosphere exchange of trace gases using the static chamber method: the precision of the instrumentation used to measure the gas concentration change in the chamber headspace, and undefined or unquantifiable sources of uncertainty. The precision-based MDF has been well studied (e.g., Parkin et al., 2012) but the undefined sources of uncertainty are less constrained and are likely related to the specific system configuration. Undefined sources of uncertainty are rarely addressed in field studies.

**Instrumental precision**

The instrumental precision for measurements of trace gas concentration changes in the chamber headspace is defined by the instrument type and maker and, for current IRGAs, it is very high and only has a minor effect on the overall uncertainty. State-of-the-art IRGAs have precisions of 0.2–1.0 ppbv for $CH_4$.

Instrumental precision-based estimation of the chamber headspace concentration change

Estimation of uncertainty based on instrumental precision has been described in detail by (Parkin et al., 2012). The uncertainty estimation is based on the accuracy of the measured trace gas concentration in the headspace of the static chamber over the incubation period. This uncertainty is dependent on the specific incubation time, number of measurements of the gas concentration in the chamber headspace, and accuracy of the instrument. The precision of the instrument determines the ability to measure a change in concentration of the gas of interest in the closure/incubation time.

Undefined sources of uncertainty

Undefined sources of uncertainty probably reflect the whole-system configuration and local changes in wind speed, pressure, and temperature. This uncertainty is difficult to estimate. One possible way to estimate this uncertainty is to set up a chamber in the field without soil contact. The chamber must be deployed with the other chambers for a sufficiently long time to characterize the undefined sources of uncertainty.

Uncertainty assessment and measurements in the field

The flux calculation from the automatic chamber system is based on the rate of concentration increase in the chamber headspace. For estimation of the instrument-based uncertainty, we used the measured concentration changes in a chamber headspace of a dummy chamber deployed in the field without any soil contact, which was automatically opened and closed using the Eosence multiplexer (eosMX (-P), Eosense Inc.; Dartmouth, Nova Scotia, Canada). During the closure (incubation) time of 300 s, measurements of the gas concentrations were taken every 10 s, except for the first 30 s. All measured gas concentrations in the headspace of the dummy chamber were used to calculate the mean and coefficient of variation. We used Monte Carlo simulations to generate 10,000 virtual chamber incubations with randomly distributed gas concentrations from those measured in the dummy chamber. Slopes from the randomly changing concentrations within the incubation time or the MDS were calculated for both summer and winter periods with significance level of $p < 0.05$ (Table S2). Calculated MDS of 0.0045 and 0.0055 ppbv $CH_4$ $s^{-1}$ for summer and winter months, respectively were lower than 0.019 nmol $m^{-2}$ $s^{-1}$ published by Christiansen et al., (2015) and similar to 0.002 nmol $m^{-2}$ $s^{-1}$ estimated by Nickerson, (2016). These estimated MDS were then removed from the calculated slopes and all slopes smaller than or equal to the MDS were assumed to be zero (or undefined). Based on the Monte Carlo simulation, increasing the number of gas concentration measurements within the incubation time will reduce the MDS. However, the MDS does not represent the total uncertainty of the *in-situ* flux measurements.

To estimate MDF that influenced by undefined sources of uncertainty during measurements by automatic chambers coupled to IRGAs, field deployment of a dummy chamber is necessary. A dummy chamber placed in the field without direct soil contact will experience the other field factors. When such a dummy chamber is deployed, an erroneous 'flux' may be detected, and this 'flux' incorporates all possible sources of uncertainty. An important advantage of using a dummy chamber is the large number of measurements possible across changes in local relative humidity, temperature, and pressure.

Workflow for estimation of the minimal detectable flux

We established six automatic soil chambers in bare and tilled field sites between 2 May and 27 July 2023 and between 26 November 2023 and 3 March 2024,. The chambers were removed from the field for 21 days in June. During these days' chambers were placed on the mesh platform and were measuring "soil fluxes" without any connection with soil. This was done to rule out the possibility that closure and opening of the chambers may affect flux estimation. One additional chamber was set on a metal sheet in the same area, without direct soil contact. All chambers were used to measure the soil $CH_4$ fluxes. The system included seven automatic chambers connected in circuit to a multiplexer by 100 m of shielded Teflon tubes. The multiplexer was connected to the ICOS in addition to a pump (KNF; 10 L $min^{-1}$; Neuberger GmbH; Freiburg, Germany) to allow the rapid recirculation of gas through the chamber–analyzer system and to avoid air-flow lag times. The multiplexer and gas analyzers were placed inside an air-conditioned shed. The chambers were measured in sequential switching mode, one after another, with 5 mins of closure time (i.e., incubation time) and 5 mins of flushing time. Each chamber measured the soil flux ~12 times per 24 h for a total of 3904 incubations during the field deployment in the rainless summer period and 6444 incubations during the winter period.

The first step in the estimation of the MDF based on the dummy chamber was the removal of outliers and exclusion of the MDS (Table S3) from the measured fluxes. The outliers were defined as slopes larger or smaller than three standard deviations of the average of all measured slopes. This was undertaken to remove potential spikes related to electrical or instrumental malfunctioning, because we did not expect any abrupt changes in point-emissions of trace gases. Removal of the MDS and outliers resulted in ~12 % of all measurements being excluded (Table S3).

After removal of the MDS and outliers, "fluxes" from the dummy chamber could be detected and attributed to undefined sources of uncertainty. A total of 664 and 1120 flux measurements from the dummy chamber were used to calculate MDF values of 0.1188 ± 0.005 and 0.0584 ± 0.0002 nmol $CH_4$ $m^{-2}$ $s^{-1}$ for summer and winter months, respectively. These, measured in field MDFs are up to 60 folds higher than reported previously MDF that were based on instrumental precision only. This approach allowed us to calculate average summer and winter fluxes (Fig. S2).

### Figures and Tables

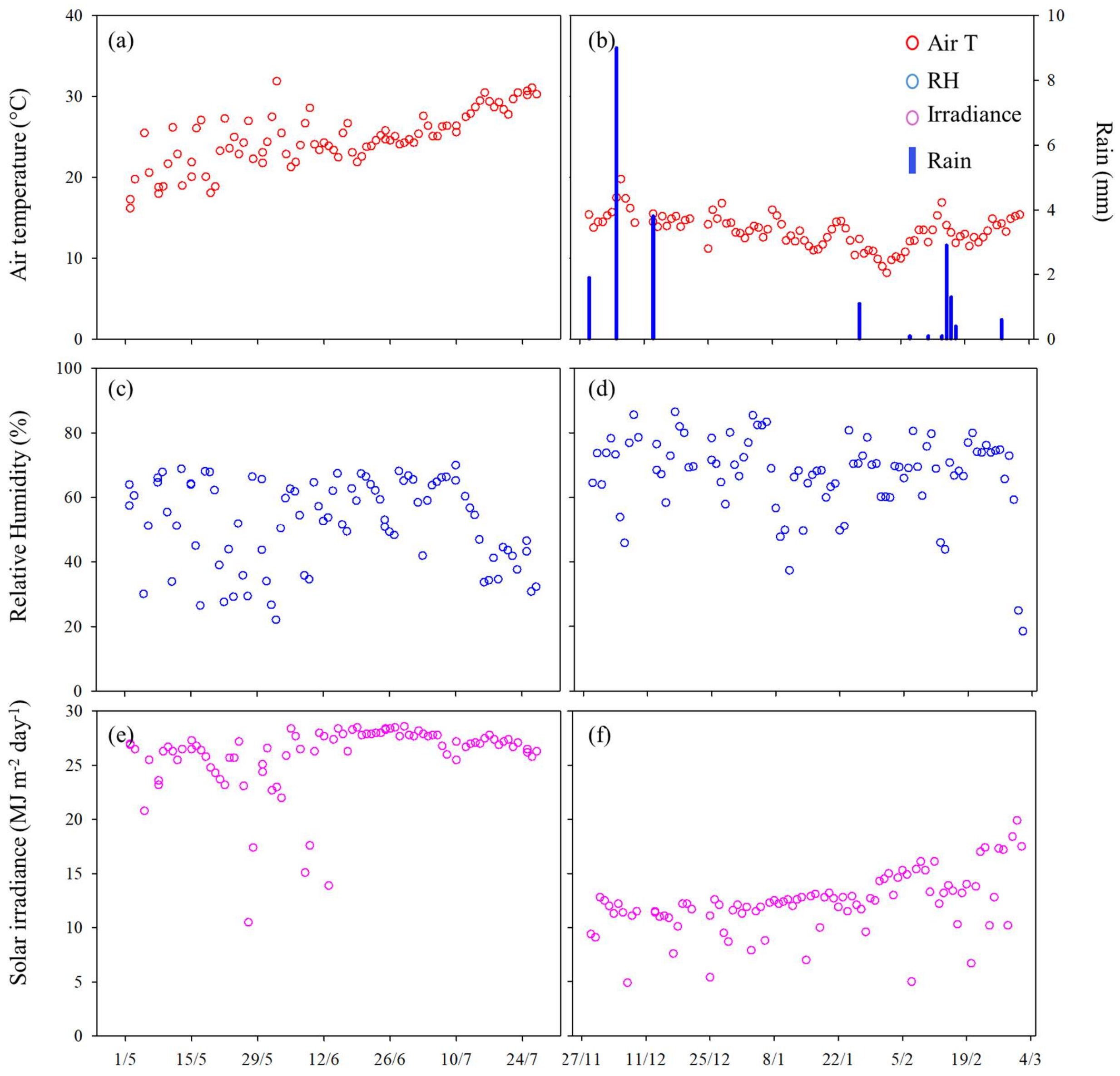


Figure S1. Climatic conditions during the experiment. Average air temperature (°C), Average relative humidity (%), and average solar irradiance (MJ $m^{-2}$ $day^{-1}$) during Summer (a, c, e) and Winter (b, d, f) months. Rainfall indicated by blue bars on panel (b).

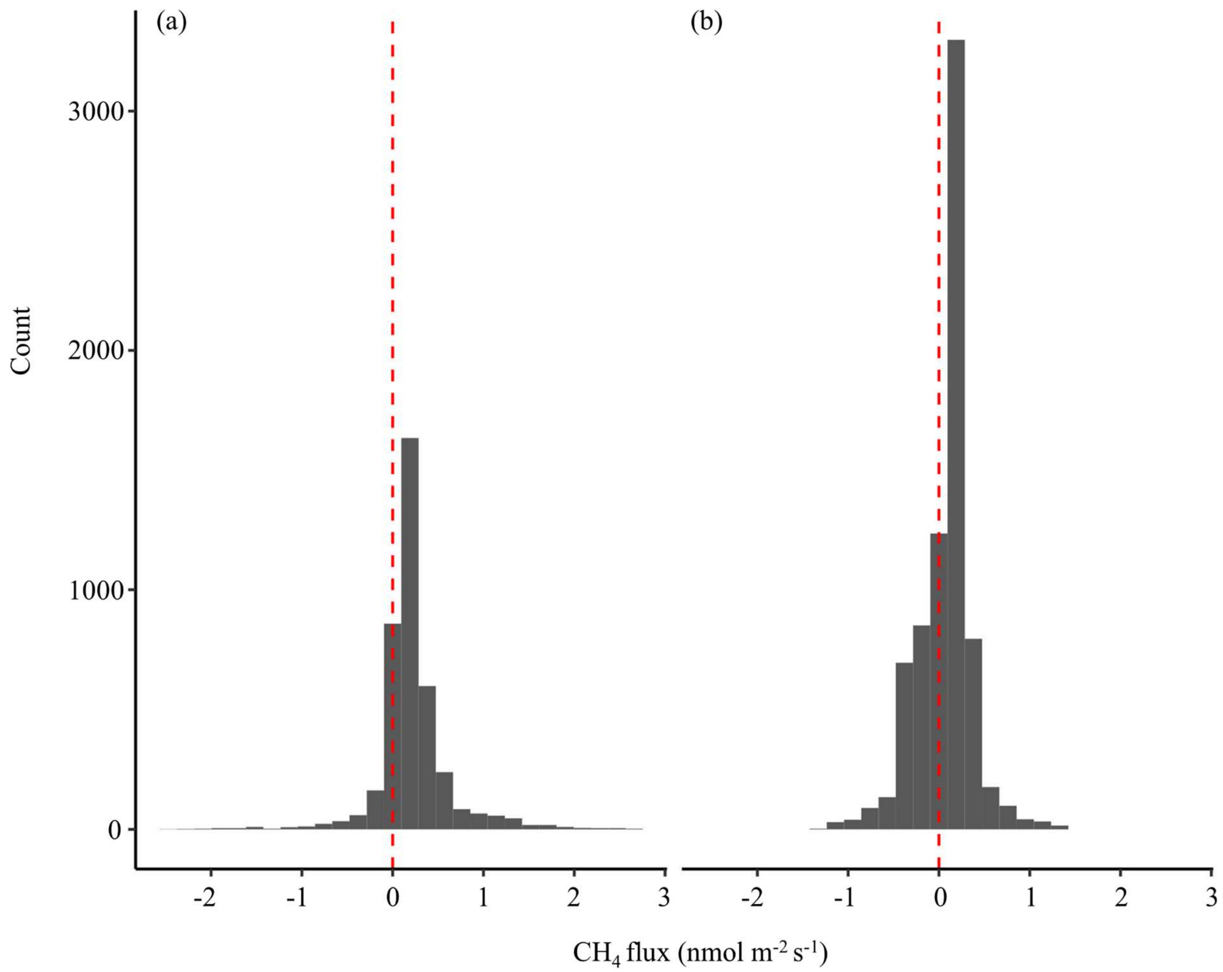


**Figure S2.** Distribution of measured soil $CH_4$ fluxes during summer (a) and winter (b) months.

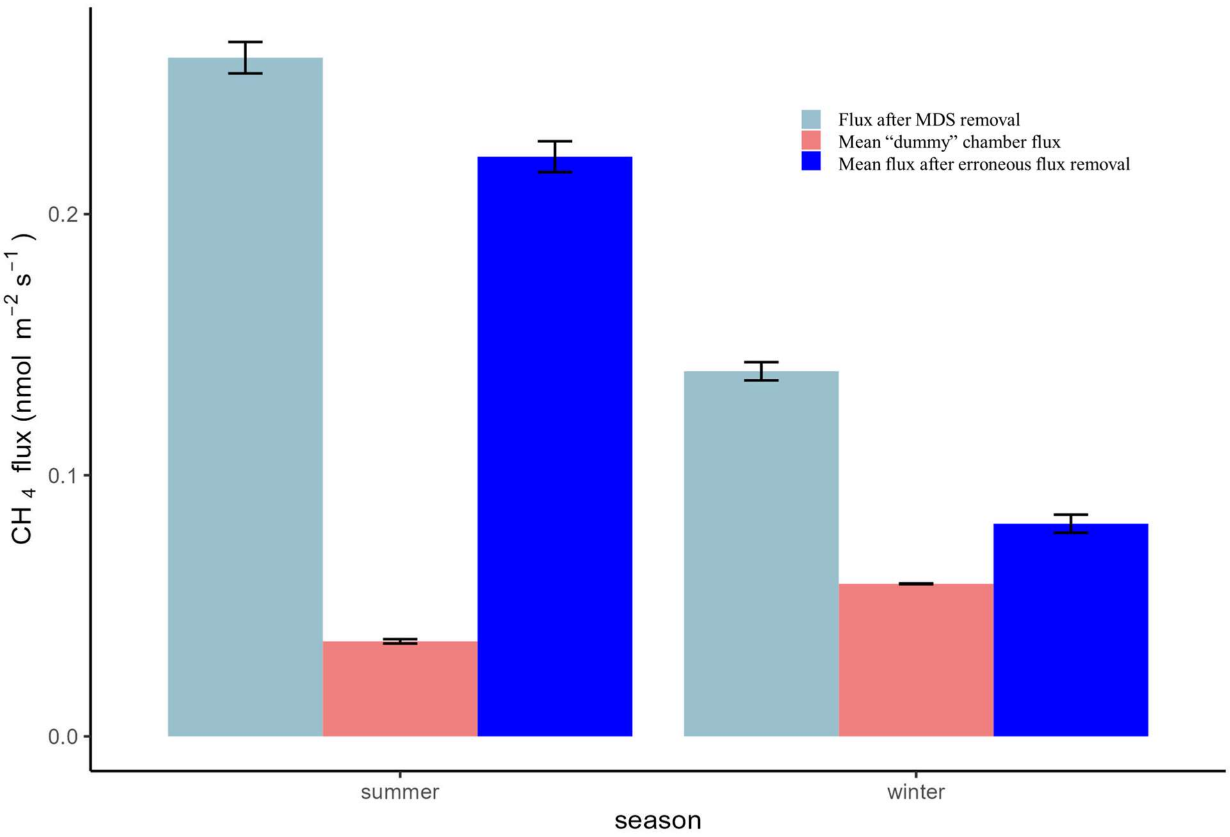


**Figure S3**. Sequential removal of the MDS and the “dummy” chamber flux during estimation of mean soil $CH_4$ flux.

**Table S1.** Descriptive statistical data for the measured $CH_4$ fluxes.

| Season | Mean | Min | Max | s.e. | Skewness | Number of measured fluxes |
|---|---|---|---|---|---|---|
| | Flux *(nmol $m^{-2}$ $s^{-1}$)* | | | | | |
| Summer | 0.222 | -2.380 | 2.696 | 0.007 | 0.517 | 3904 |
| Sun | 0.296 | -0.34 | 0.78 | 0.005 | 0.193 | 994 |
| Shade | 0.144 | -0.38 | 0.77 | 0.005 | 0.046 | 999 |
| Winter | 0.081 | -1.246 | 1.362 | 0.004 | -0.397 | 6444 |

**Table S2.** Minimum detectable slopes of the gaseous concentration change in an automatic chamber deployed in the field on a metal sheet and detached from the soil (the dummy chamber). The calculation of the MDS is based on the 17,928 – 30,240 measured concentrations during 653 and 1120 individual incubations of the dummy chamber in the summer and winter months, respectively.

| *Season* | $CH_4$ *ppbv sec$^{-1}$* |
|---|---|
| *Summer* | ±0.0045 |
| *Winter* | ±0.0055 |

**Table S3.** Proportion of incubations removed from the analysis due to the minimum detectable slope (MDS) and outliers' removal.

| | $CH_4$ | |
|---|---|---|
| | % | |
| | *Outliers* | MDS |
| *Summer* | 1.46 | 6.73 |
| *Winter* | 1.64 | 1.81 |

**Table S4.** Parameters are used to identify suitable areas for estimation of dry soil $CH_4$ efflux, globally.

| Parameter | Value | Unit |
|---|---|---|
| Bulk Density | 120 – 170 | kg×10 $m^{-3}$ |
| Clay | 5 – 20 | % |
| Sand | 10 – 90 | % |
| pH | 7 – 11 | - |
| Precipitation | 0 – 210 | mm |
| Precipitation in warmest months | 0 – 5 | mm |

Table S5. ANOVA of soil carbon content effect on $CH_4$ fluxes during summer and winter months. Shown are numerator (Num DF) and denominator (Den DF) degrees of freedom, F-values and p-values for the Statistically significant effects are indicated by **bold** ($P < 0.05$). The data was subjected to log (summer) and exponential (winter) transformation prior to the analysis.

| Effect | Num DF | Den DF | F value | Pr>F |
|---|---|---|---|---|
| Summer | | | | |
| Carbon content | 3 | 10.73 | 4.84 | **0.0226** |
| Winter | | | | |
| Carbon content | 3 | 9.27 | 10.32 | **0.0026** |
